# Assessing the perceived reverberation in different rooms for a set of musical instrument sounds

**DOI:** 10.1101/2020.03.13.984542

**Authors:** Alejandro Osses Vecchi, Glen McLachlan, Armin Kohlrausch

## Abstract

Previous research has shown that the perceived reverberation in a room, or reverberance, depends on the sound source that is being listened to. In a study by Osses et al. [(2017) J. Acoust. Soc. Am. **141**(4), EL381-EL387], reverberance estimates obtained from an auditory model for 23 musical instrument sounds in 8 rooms supported this sound-source dependency. As a follow-up to that study, a listening experiment with 24 participants was conducted using a subset of the original sounds with the purpose of mapping each test sound onto a reverberance scale. The experimentally-obtained reverberance estimates were significantly correlated with the simulated reverberance, providing further evidence that the sensation of reverberance is sound-source dependent.

## 1. Introduction

For the acoustic characterization of a room, metrics that are derived from measured room impulse responses (RIRs) such as reverberation time (RT) and clarity are often used. An implicit assumption of this practice is that rooms can be characterized as linear time-invariant (LTI) systems (e.g., Vorländer, 2008, Chapter 7) or, in other words, that the interplay between sound source and room is independent of the sound source and receiver. When (human) listeners are the end users of a room (the “receivers”) the LTI assumption is unlikely to be valid given that the hearing system introduces a non-linear sound processing that varies over time due to factors such as attention (Alain, 2007) or aging (Moore, 2019). If the LTI assumption fails then impulsive sounds will in general not be representative of actual listening experiences in a room.

Evidence against the LTI assumption is given by Teret et al. (2017), who asked 8 participants to match the perceived RT of subsequent signal pairs using either identical or different signals with varying degrees of reverberation. They found that the participants’ responses were more accurate when the task was conducted with identical signals. From this the authors inferred that the just-noticeable difference (JND) in RT from other studies may be valid when listening to identical signals only. This hypothesis is also supported by Klockgether and van de Par (2016), whose experimental results show that the JNDs for several metrics depend on the nature of the sound source under test. They observed that the JND of a percussive sound was more robust to reverberation than tonal sounds, with lower and higher JNDs, respectively.

In the same line of reasoning but with the specific goal of testing the dependency of reverberance on sound source type, Osses et al. (2017) used an artificial listener based on the Room Acoustic Analyzer model (RAA, van Dorp et al., 2013), to assess the reverberance of a set of orchestra sounds auralized in 8 different rooms. Their results showed that the model estimates depended on the spectral properties and the presentation level of the sounds, despite the fact that average estimates were significantly correlated with the RIR-derived RT estimates of *T*_30_ and early decay time (EDT). To provide experimental evidence for those model predictions the present study used a subset of the same orchestra sounds and rooms and a listening experiment was conducted. Our expectations were that: (1) experimental reverberance estimates (*P* _REV,exp_) are instrument-dependent, and (2) *P* _REV,exp_ estimates are correlated with simulated reverberance estimates (*P* _REV,sim_) obtained using the RAA model.

## 2. Method

### 2.1. Stimuli

Our stimuli consisted of anechoic recordings from 8 instruments playing the 3^rd^ movement of Brahms Symphony Nr. 4 (Rindel, 2015) that were auralized using the binaural RIRs of 8 different rooms. These sounds are a subset of the stimuli used by Osses et al. (2017) and we chose them to be representative of their published simulation results. Using 90-s-long sounds they found that simulated reverberance estimates for different instruments (labeled here as *P* _REV,sim,90 s_) followed three different reverberant trends: trend 1, 2, and 3 (“other trends”). From trend 1 we chose: violin, French horn, and timpani. From trend 2 we chose: flute, piccolo, and trumpet. From trend 3 we chose double bass and contrabassoon. We reduced the duration of the excerpts to no more than 9 s. More specifically, we used the excerpts corresponding to bars 10 to 18 of the symphony, where all instruments play *fortissimo*.

The reverberant orchestra sounds were obtained by digital convolution of the 8 selected anechoic recordings using the binaural RIRs of 8 rooms (see Table 1). The convolution was performed in MATLAB, where a fixed gain of −9 dB was applied to avoid clipping while storing the waveforms. The auralized sounds were calibrated to produce a sound pressure level (SPL) of 107.7 dB at the maximum possible amplitude (0 dBFS= 107.7 dB SPL). The resulting sounds had levels that we labeled as comfortable (see Table 2).

**Table 1.** List of rooms used in this study sorted by increasing EDT times. Column G gives an indication of the sound strength in the rooms. An extended version of this table can be found in (Osses et al., 2017, their Table 2).

| Room | Volume (m<sup>3</sup>) | EDT (s) | T<sub>30</sub> (s) | G (dB) | Room | Volume (m<sup>3</sup>) | EDT (s) | T<sub>30</sub> (s) | G (dB) |
| --- | --- | --- | --- | --- | --- | --- | --- | --- | --- |
| 1 | 14000 | 0.80 | 1.14 | 5.4 | 5 | 23000 | 1.24 | 2.01 | 1.3 |
| 2 | 23000 | 0.81 | 1.20 | 0.0 | 6 | 2500 | 1.27 | 1.34 | 10.8 |
| 3 | 14000 | 0.83 | 1.51 | 7.3 | 7 | 15000 | 1.47 | 2.23 | 8.5 |
| 4 | 2500 | 1.04 | 1.16 | 9.8 | 8 | 2500 | 2.48 | 2.51 | 12.7 |

**Table 2.**
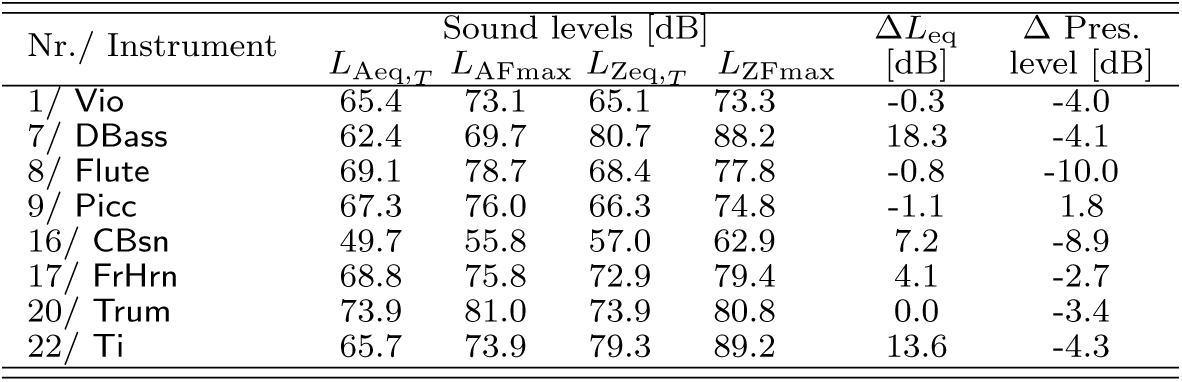
Level information (average across rooms) of the sounds used in the listening experiment. The levels 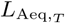 (A-weighted) and 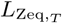 were integrated over the entire duration (9 s) and their difference is indicated as Δ*L*_eq_. The column “Δ Pres. level” was obtained as the difference between the maximum value of the auralized waveforms *L*_AFmax_ and the maximum of the 90-s sounds (Osses et al., 2017, *L*_AFmax_ in their Table 3). The overall differences are negative, meaning that similar (Picc) or softer reproduction levels were used in our listening experiments with respect to the sounds as used by Osses et al. (2017).

| Nr./ Instrument | Sound levels [dB] | | | | $\Delta L_{\text{eq}}$ [dB] | $\Delta$ Pres. level [dB] |
| --- | --- | --- | --- | --- | --- | --- |
| | $L_{\text{Aeq},T}$ | $L_{\text{AFmax}}$ | $L_{\text{Zeq},T}$ | $L_{\text{ZFmax}}$ | | |
| 1/ Vio | 65.4 | 73.1 | 65.1 | 73.3 | -0.3 | -4.0 |
| 7/ DBass | 62.4 | 69.7 | 80.7 | 88.2 | 18.3 | -4.1 |
| 8/ Flute | 69.1 | 78.7 | 68.4 | 77.8 | -0.8 | -10.0 |
| 9/ Picc | 67.3 | 76.0 | 66.3 | 74.8 | -1.1 | 1.8 |
| 16/ CBsn | 49.7 | 55.8 | 57.0 | 62.9 | 7.2 | -8.9 |
| 17/ FrHrn | 68.8 | 75.8 | 72.9 | 79.4 | 4.1 | -2.7 |
| 20/ Trum | 73.9 | 81.0 | 73.9 | 80.8 | 0.0 | -3.4 |
| 22/ Ti | 65.7 | 73.9 | 79.3 | 89.2 | 13.6 | -4.3 |

### 2.2. Apparatus

The stimuli were presented binaurally via Sennheiser HD 265 Linear circumaural headphones (Sennheiser, Wedemark, Germany), while the participants were seated in a single-walled sound booth. The data were collected using the software Web Audio Evaluation (Jillings et al., 2016) using Google Chrome on a local computer.

### 2.3. Participants and power analysis

Twenty-four participants (5 females, 19 males) were recruited from the JF Schouten subject database of the TU/e university. At the time of testing the participants were between 19 and 43 years old (average of 24 years) and they all had self-reported normal hearing. They provided their informed consent before starting the experimental session and were paid for their contribution.

The experiment used a repeated measures (within-subject) design with interest to check the main effects of the factors “musical instrument” and “room.” Sixty-four sound stimuli were used, which could be grouped into either 8 groups of 8 instruments or 8 groups of 8 rooms. The first case was of more interest to us, with a null-hypothesis that reverberance estimates are the same for the 8 instruments. Assuming a medium effect size (Cohen’s *f* = 0.25), and *α* level of 0.05, 24 participants had to be recruited to support/reject the hypothesis with a power *β* ≥0.90 (*β*_actual_ =0.96).

### 2.4. Experimental procedure

A multi-stimulus comparison method was used (De Man and Reiss, 2013), where participants were asked to sort 8 stimuli along a continuous scale ranging from 0 to 1 according to an increasing sensation of reverberance. Sixteen trials (16 scales with 8 stimuli each) were presented to each participant, with 8 trials having stimuli of the same instrument in different rooms (within-instrument), and 8 trials having different instruments in the same room (within-room). All trials were completed within a single one-hour session.

### 2.5. Simulation data: Predicted reverberance P_REV_

Given that the presentation levels of the 9-s excerpts used in our study differ from the levels used in the previously published simulations, we reassessed the reverberance estimates using a custom MATLAB implementation^1^ (Osses, 2017; Osses et al., 2017) of the RAA model (van Dorp et al., 2013), where reverberance estimates were obtained for 5-s long sections with an 80% overlap, resulting in five *P* _REV,sim_ estimates for each stimulus. The obtained average estimates are shown in Fig. 1.

**Fig. 1.**
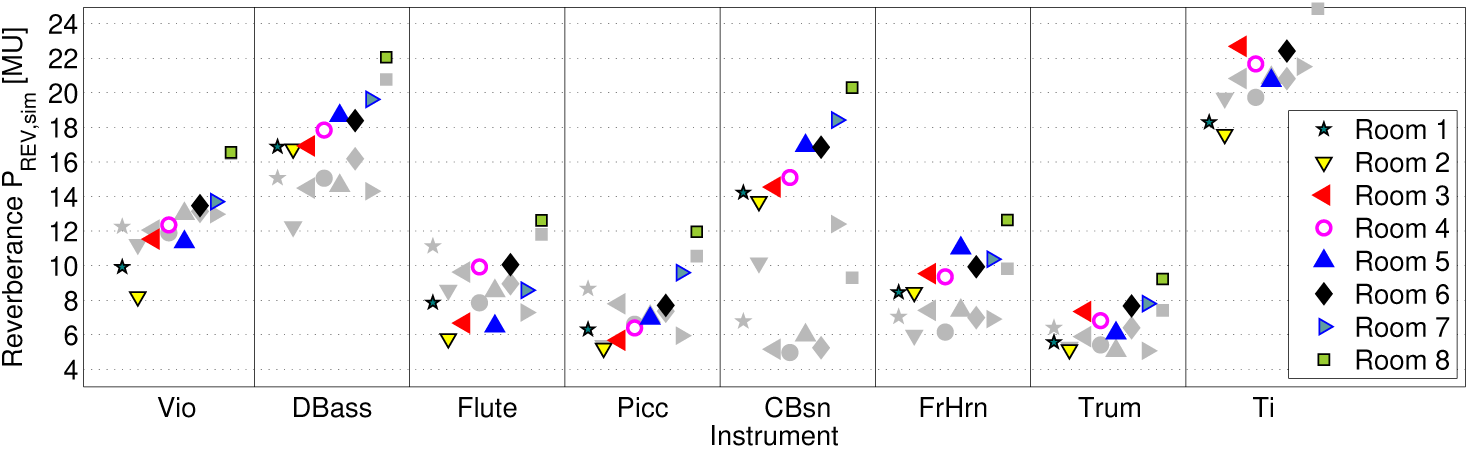
Simulated *P* _REV,sim_ estimates expressed in model arbitrary units (MU) for the eight selected musical instruments in the eight acoustic environments (Rooms 1–8). For ease of comparison, the simulated *P*_REV,sim 90 s_ estimates reported by Osses et al. (2017) are indicated by gray markers.

## 3. Results

The results for the within-instrument evaluations are shown in Fig. 2A, where the median reverberance estimates *P* _REV,exp_ ranged between 0.10 (`Vio` in Room 2) and 0.98 (`DBass` in Room 8). Since the sounds were compared within instruments, the individual scales may not be directly related to each other. This is because the participants’ responses only required to be referenced to the sound samples within each trial.

**Fig. 2.**
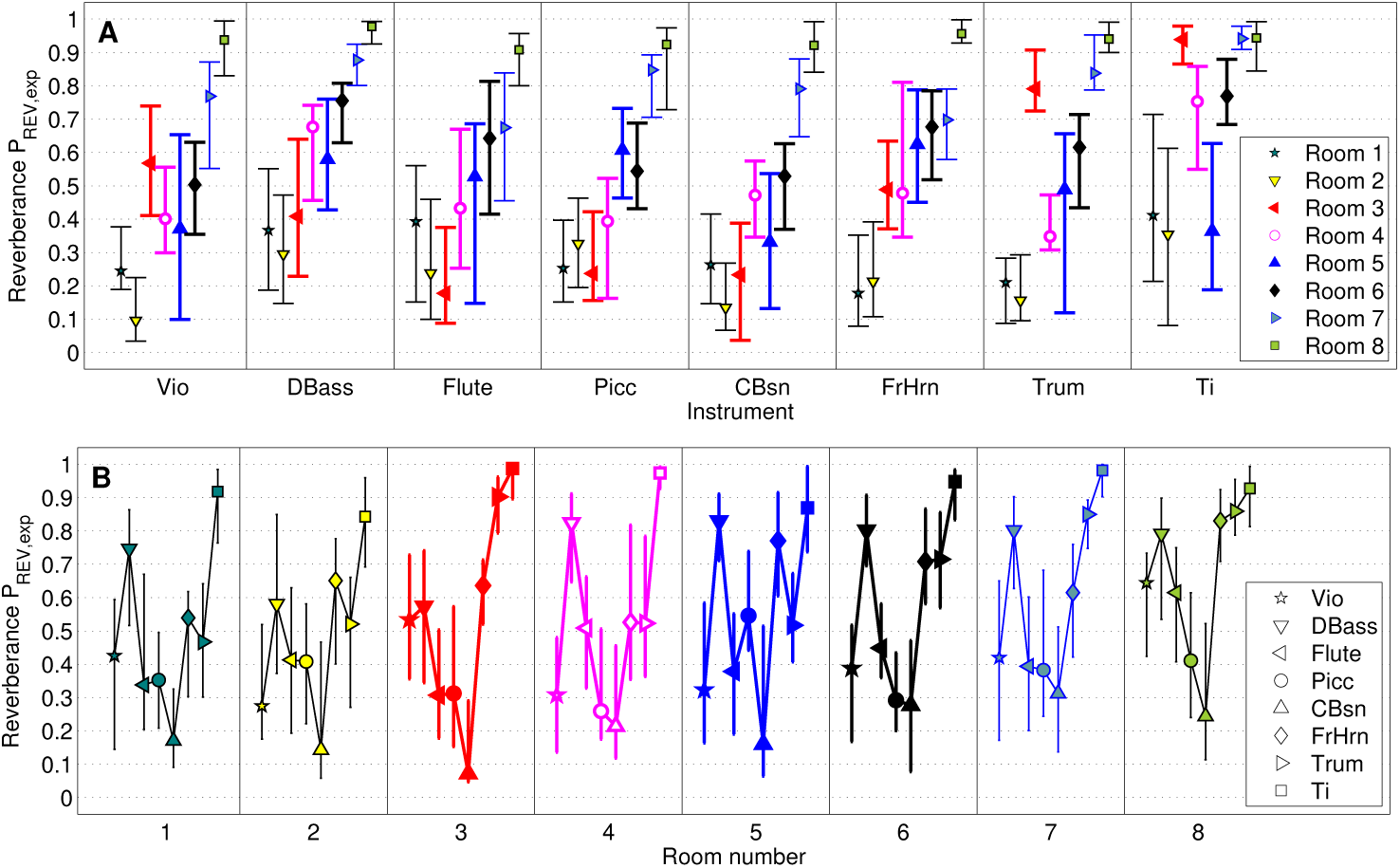
Experimental results from the listening test in the (A) within-instrument and (B) within-room conditions. The median reverberance estimates *P* _REV,exp_ in 8 different acoustic environments are indicated together with their corresponding interquartile ranges.

The results for the within-room evaluations are shown in Fig. 2B. The median reverberance estimates ranged between 0.07 (`CBsn` in Room 3) and 0.99 (`Ti` in Room 3). The instruments sorted in order of increasing reverberance were: `CBsn` (P_REV,exp_= 0.20), `Picc`, `Vio`, `Flute` (P_REV,exp_= 0.37 ≈ 0.41 ≈ 0.43), `FrHrn`, `Trum` (P_REV,exp_= 0.66 ≈ 0.67), `DBass` (P_REV,exp_= 0.74), and `Ti` (P_REV,exp_= 0.93).

## 4. Discussion

### 4.1. Within-instrument evaluation

#### Comparison with the simulation data

The experimental data *P* _REV,exp_ (Fig. 2) were compared with average simulated *P* _REV,sim_ estimates obtained from our 9-s long stimuli and with the previously reported average and maximum simulated estimates (*P* _REV,sim,90 s_ and *P* _REV,sim,90 s,max_). The corresponding (Pearson) correlation values are shown in Table 3. *P* _REV,exp_ was significantly correlated with *P* _REV,sim_ with *r*_*p*_ values between 0.81 (`Flute`) and 0.94 (`Vio` and `FrHrn`). When comparing *P* _REV,exp_ with *P* _REV,sim,90 s_ only three correlation values were significant (`Vio`, `DBass`, `FrHrn`) and one approached significance (`Ti`). Although in this case *r*_*p*_ values were expected to be lower because the *P* _REV,sim,90 s_ estimation considered sound parts that were never presented to our listeners, these estimates could be interpreted as belonging to a more representative playing context of the instruments. Since the selected instruments played fortissimo during the 9-s excerpts, the correlation between *P* _REV,exp_ and *P* _REV,sim,90 s,max_ was included.^2^ In this case, six (of eight) *r*_*p*_ values were significant with values between 0.74 (`Trum` and `Ti`) and 0.91 (`DBass`).

**Table 3.** Pearson correlation *r*_*p*_ between experimental and simulated *P* _REV_ estimates in the within-instrument condition. Each *r*_*p*_ value is obtained by comparing 8 pairs of data points (6 degrees of freedom).

| Nr./ Instrument | $P_{\text{REV,exp}}$ correlated with | | |
| --- | --- | --- | --- |
| | $P_{\text{REV,9 s}}$ | $P_{\text{REV,90 s}}$ | $P_{\text{REV,max,90 s}}$ |
| 1/ Vio | 0.94* | 0.81* | 0.77* |
| 7/ DBass | 0.90* | 0.72* | 0.91* |
| 8/ Flute | 0.81* | 0.22 | 0.46 |
| 9/ Picc | 0.93* | 0.27 | 0.26 |
| 16/ CBsn | 0.93* | 0.42 | 0.77* |
| 17/ FrHrn | 0.94* | 0.73* | 0.90* |
| 20/ Trum | 0.92* | 0.35 | 0.74* |
| 22/ Ti | 0.88* | 0.62** | 0.74* |

(\*) Significant correlation, $p < 0.05$ . (\*\*) Correlations that approach significance, $p < 0.10$ .

#### Existence of reverberance trends

To investigate the existence of the reverberance trends that had been previously observed (Osses et al., 2017), the *P* _REV,exp_ estimates of Fig. 2 were first used to construct the correlation (“similarity”) matrix shown in Table 4, which was subsequently subjected to multidimensional scaling (MDS) to derive a graphical two-dimensional representation of the 8 test instruments. The resulting Cartesian representation is shown in Fig. 3^3^. From the figure, the following can be observed: (1) the instruments from trend 1 (black markers: `Vio`, `FrHrn`, `Ti`) are placed in the neighborhood of each other; (2) In trend 2 (red markers), `Flute` and `Picc` are close to each other, with `Trum` being located slightly farther apart and getting somewhat closer to `FrHrn` from trend 1, and; (3) In trend 3 (white markers) both instruments are very close to each other. In other words, based on this Cartesian representation, each instrument can be grouped into a class (trend) defined by one of the three nonlinearly separable areas (dashed lines in Fig. 3). These results, however, should be interpreted with care given that this representation was obtained with a lower number of stimuli (8 instruments) than used in the trend analysis by Osses et al. (2017) (14 instrument groups). This implicitly assumes that (1) none of the omitted instruments would significantly affect the position of the 8 instruments in the obtained space, and that (2) the correlation matrix of Table 4 is a faithful representation of the similarity among musical instruments.

**Table 4.**
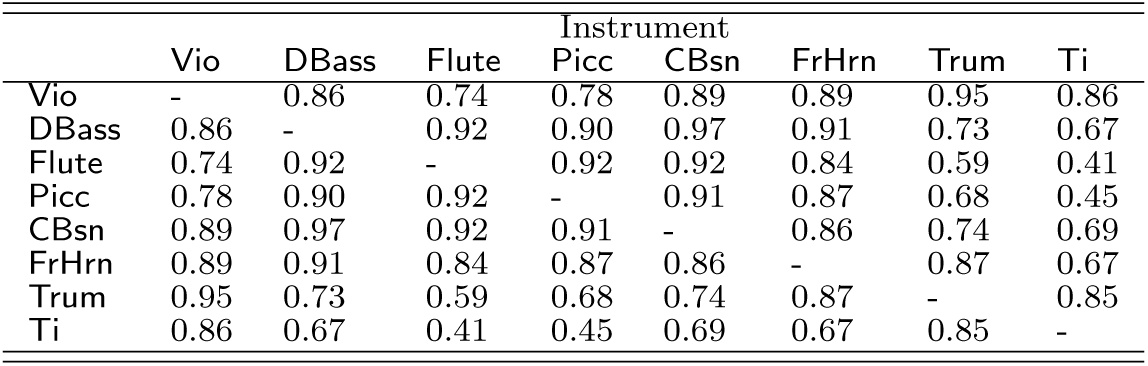
Pearson correlation matrix among experimental estimates *P* _REV,exp_ for all possible instrument pairs. The lowest and highest correlation values were found for the comparisons between flute and timpani estimates (*r*_*p*_ = 0.41) and between double bass and contrabassoon estimates (*r*_*p*_ = 0.97), respectively.

|  | Instrument |  |  |  |  |  |  |  |
| --- | --- | --- | --- | --- | --- | --- | --- | --- |
|  | Vio | DBass | Flute | Picc | CBsn | FrHrn | Trum | Ti |
| Vio | - | 0.86 | 0.74 | 0.78 | 0.89 | 0.89 | 0.95 | 0.86 |
| DBass | 0.86 | - | 0.92 | 0.90 | 0.97 | 0.91 | 0.73 | 0.67 |
| Flute | 0.74 | 0.92 | - | 0.92 | 0.92 | 0.84 | 0.59 | 0.41 |
| Picc | 0.78 | 0.90 | 0.92 | - | 0.91 | 0.87 | 0.68 | 0.45 |
| CBsn | 0.89 | 0.97 | 0.92 | 0.91 | - | 0.86 | 0.74 | 0.69 |
| FrHrn | 0.89 | 0.91 | 0.84 | 0.87 | 0.86 | - | 0.87 | 0.67 |
| Trum | 0.95 | 0.73 | 0.59 | 0.68 | 0.74 | 0.87 | - | 0.85 |
| Ti | 0.86 | 0.67 | 0.41 | 0.45 | 0.69 | 0.67 | 0.85 | - |

**Fig. 3.**
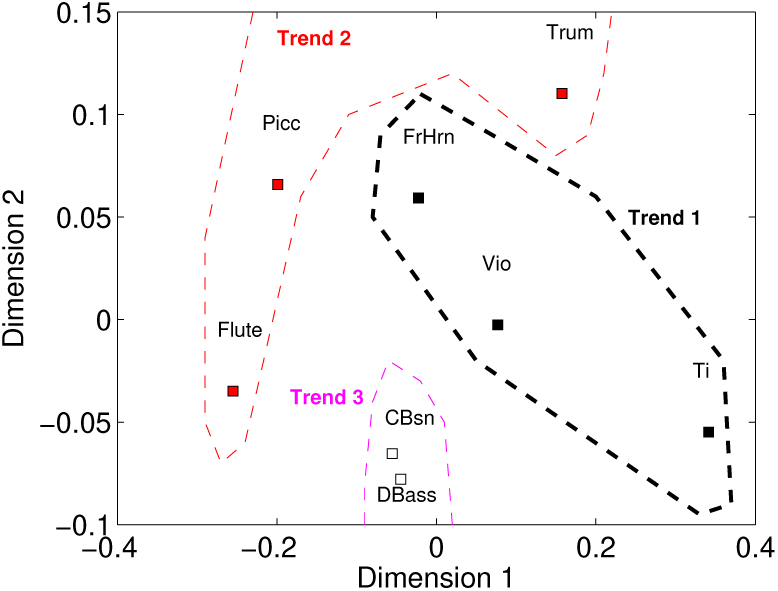
Graphical representation of the 8 orchestra instruments used in the listening experiment. This analysis is based on the correlation matrix of *P* _REV,exp_ estimates that is shown in Table 4. The colors and dashed lines indicate the three trends that were previously found using simulated reverberance estimates (Osses et al., 2017) with black, red, and white markers for trends 1, 2, and 3, respectively.

### 4.2. Sound source dependency: Within-room evaluation

The within-room results shown in Fig. 2B can be directly used to evaluate the dependency of reverberance on the sound source type. For Room 8, which is the most reverberant room in our dataset, the instruments sorted from low to high scores, i.e., from least to most reverberant were: `CBsn`, `Picc`, `Flute`, `Vio`, `DBass`, `FrHrn`, `Trum`, and `Ti`, respectively. This “reverberance pattern” is similar in the other seven acoustic environments, with a rank-order (Spearman) correlation that ranges between 0.69 (*r*_*s*_ for Room 5) and 0.98 (*r*_*s*_ for Room 3).

The average *P* _REV,exp_ estimates between 0.20 (`CBsn`) and 0.93 (`Ti`) may be used as evidence for the reverberance dependency on the sound source (instrument) type. To provide further statistical evidence, a repeated measures one-way ANOVA (one for each room) was conducted to analyze the influence of the variable “instrument” on *P* _REV,exp_. The results show that “instrument type” influenced significantly the experimental reverberance scores in all rooms, as shown in Table 5.

**Table 5.** Results of repeated measures one-way ANOVAs conducted for each acoustic environment. In all the analyses it was found that the variable “instrument” influences significantly the experimental *P* _REV,exp_ values obtained in the within-room evaluations. For each of the eight acoustic environments, 192 observations were available (8 instruments evaluated once by 24 participants).

| Room | $F(7, 184)$ | $p$ | Room | $F(7, 184)$ | $p$ |
| --- | --- | --- | --- | --- | --- |
| 1 | 34.82 | < 0.001 | 5 | 16.91 | < 0.001 |
| 2 | 15.52 | < 0.001 | 6 | 25.96 | < 0.001 |
| 3 | 19.61 | < 0.001 | 7 | 27.94 | < 0.001 |
| 4 | 10.55 | < 0.001 | 8 | 22.93 | < 0.001 |

## 5. Conclusions

This study was focused on the experimental validation of previously published reverberance predictions (Osses et al., 2017) by conducting a listening experiment using similar sound stimuli and the same room acoustic environments. We adopted a multi-stimulus comparison method to estimate the reverberance of 64 stimuli: 8 musical instrument sounds auralized in 8 different rooms (EDT between 0.80 and 2.48 s). The sounds were presented in blocks of 8 stimuli in either of two conditions: within-instrument or within-room condition. The results of the within-instrument conditions showed that (1) simulated reverberance estimates (*P* _REV,sim_, *P* _REV,sim,90 s_) obtained using the binaural auditory model by van Dorp et al. (2013), as implemented by Osses et al. (2017), are significantly correlated with the collected experimental data; and that (2) the three trends obtained in our previous study could also be observed if instruments with similar *P* _REV,exp_ are grouped together. We indicated, however, that this latter analysis may require further validation. In turn, the results of the within-room conditions supported the significant influence of instrument type on reverberance for all tested rooms.

Two aspects that require further attention are the extent to which reverberance depends on sound presentation level and spectral content of the stimuli, both aspects that have already been identified as relevant (Osses et al., 2017; Lee et al., 2012). In our experimental approach, the spectral dependency was only indirectly accounted for in our trend analysis, whereas the sound presentation level was not varied, accounting only for natural level differences between instruments and rooms. The investigation of these aspects requires further experimental data collection.

In a broader context, this paper showed that perception-based predictions of a room acoustic indicator, in this case of reverberance (see also Lee et al., 2017), can be correlated with actual listening experiences in a room that depend on both “sound source” and “receiver.” This is in contrast to established measurement guidelines (ISO 3382-1, 2009), where the acoustic properties of a room are considered as linear and time invariant, that is, room properties are assumed to be independent of the type of excitation, and of the level of the exciting signals. Such a source-filter characterization allows to characterize rooms as LTI systems. The results of this paper are in line with previous findings (Klockgether and van de Par, 2016; Teret et al., 2017) and may be used as evidence that the perception of room acoustic parameters depends on the context for which the room is used and this is contrary to the idea of an LTI system.

## Acknowledgments

This research work has been funded by the European Commission within the ITN Marie Skłodowska Curie Action project BATWOMAN under the 7^th^ Framework Programme (EC grant agreement N°605867).

## References and links

1. The MATLAB implementation of the RAA model (Osses, 2017) was built using the framework of the Auditory Modelling Toolbox (Soendergaard and Majdak, 2013).
2. 2The maximum simulated reverberance estimate PREV,sim,90 s,max was obtained from percentile 75 of the frame-based reverberance estimates of each 90-s long sound.
3. For details about the data processing the reader is referred to (Osses, 2018, Chapter 6).

